# Inferring decoding strategies for multiple correlated neural populations

**DOI:** 10.1101/108019

**Authors:** Kaushik J Lakshminarasimhan, Alexandre Pouget, Gregory C DeAngelis, Dora E Angelaki, Xaq Pitkow

## Abstract

Studies of neuron-behaviour correlation and causal manipulation have long been used separately to understand the neural basis of perception. Yet these approaches sometimes lead to drastically conflicting conclusions about the functional role of brain areas. Theories that focus only on choice-related neuronal activity cannot reconcile those findings without additional experiments involving large-scale recordings to measure interneuronal correlations. By expanding current theories of neural coding and incorporating results from inactivation experiments, we demonstrate here that it is possible to infer decoding weights of different brain areas without precise knowledge of the correlation structure. We apply this technique to neural data collected from two different cortical areas in macaque monkeys trained to perform a heading discrimination task. We identify two opposing decoding schemes, each consistent with data depending on the nature of correlated noise. Our theory makes specific testable predictions to distinguish these scenarios experimentally without requiring measurement of the underlying noise correlations.

**Author Summary:** The neocortex is structurally organized into distinct brain areas. The role of specific brain areas in sensory perception is typically studied using two kinds of laboratory experiments: those that measure correlations between neural activity and reported percepts, and those that inactivate a brain region and measure the resulting changes in percepts. The two types of experiments have generally been interpreted in isolation, in part because no theory has been able combine their outcomes. Here, we describe a mathematical framework that synthesizes both kinds of results, giving us a new way to assess how different brain areas contribute to perception. When we apply our framework to experiments on behaving monkeys, we discover two models that can explain the perplexing finding that one brain area can predict an animal’s percepts, even though the percepts are not affected when that brain area is inactivated. The two models ascribe dramatically different efficiencies to brain computation. We show that these two models can be distinguished by an experiment that measures correlations while inactivating different brain areas.

## Introduction

Although much is known about how single neurons encode information about stimuli, how neurons contribute to percepts is less well understood[1]. The latter, called the “decoding problem”, seeks to identify how the brain uses the information contained in neuronal activity. Although some studies have sought to understand *principled* ways to decode population responses in the presence of correlated noise [2–12], the rules by which the brain *actually* integrates information across noisy neurons remain unclear.

Neuroscientists have traditionally investigated this question using two distinct approaches: causal or correlational. In causal approaches, experimenters selectively activate or inactivate brain regions of interest, and measure resulting perceptual or behavioural changes. In correlational approaches, experimenters measure correlations between behavioural choices and neuronal activity, typically quantified by ‘choice probability’ (reviewed in Ref. [13]) or, more straightforwardly, by ‘choice correlation’ (CC)[14,15]. If CCs reflect a functional link between neurons and behaviour, one would expect brain areas with greater CCs to contribute more strongly to behaviour. This naïve view is contradicted by recent results that reveal a striking dissociation between the magnitude of CCs and the effects of inactivation across brain systems in rodents[16,17] and primates[18,19]. In hindsight, this apparent disagreement is not all that surprising because the two techniques, on their own, yield results whose interpretation is fraught with major difficulties.

For instance, the CC of a neuron depends not only on its direct influence on behaviour but also on the influence of all the other neurons with which it is correlated. As an extreme example, a neuron that is not decoded at all could be correlated with one that is, and thus exhibit choice-related activity[9]. Recent theoretical results show that it is possible, in principle, to use knowledge of noise correlations to extract decoding weights from CCs[14]. However, directly measuring the correlational structures that matter for decoding may be extremely difficult[20]. This problem is compounded by the fact that behaviourally relevant information may be distributed across neurons in multiple brain areas, so neuronal CCs in one area may depend on activity in other areas. Moreover, in causal approaches, inactivation of one brain area could lead to a dynamic recalibration of decoding weights from other areas. Therefore, changes in behavioural thresholds following inactivation may not be commensurate with the contribution of the area.

When analysed in conjunction, however, results from correlational and causal studies may together provide constraints that can be used to precisely determine the relative contributions of the brain areas involved. In this work, we extend recent theories[14,15,20] and propose a general framework for inferring decoding weights of neurons across multiple brain areas using CCs and changes in behavioural threshold following inactivation. The two quantities together provide a direct estimate of the relative contributions of different areas without needing to precisely measure the correlation structure. We demonstrate our technique by applying it to data from macaque monkeys trained to perform a heading discrimination task. In this task, there is a known discrepancy[18,21–23] between CCs and the effects of inactivating two brain areas: although neurons in the ventral intraparietal (VIP) area were found to be substantially better predictors of the animal’s choices than dorsal medial superior temporal (MSTd) neurons, performance is impaired by inactivating MSTd but not VIP. We use our framework to extract key properties of the decoder that can account for these counter-intuitive results. To our surprise, we find that, depending on the structure of correlated noise, experimental data are consistent with two opposing schemes that attribute either too much or too little weight to VIP. We use our theory to make specific testable predictions to distinguish these schemes using CCs measured during inactivation, again without measuring the detailed noise correlations.

## Results

### Decoding framework

We consider a linear feedforward network in which the firing rates **r** of the neurons are combined linearly using weights **w** to yield a locally unbiased estimate 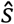 of the stimulus according to 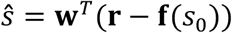, where **f**(*s*_0_) is the mean response to a reference stimulus *s*_0_. In each trial, the animal is assumed to reach a binary decision given by 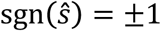, where sgn is the signum function. For a decoder that linearly reads out neurons from two subpopulations, *x* and *y*, the estimate 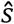 can be expressed as:

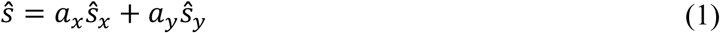

where 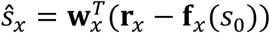 and 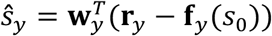 denote unbiased estimates derived from neurons in subpopulations *x* and *y* respectively. Thus the problem of decoding multiple populations can be viewed as one of scaling and combining estimates from individual populations. Note that this is equivalent to a single linear decoder of both populations together using w = [*a_x_***w**_*x*_ *a*_*y*_**w**_*y*_] The form of equation (1) has two advantages: (*i*) it is easy to identify and compare the relative contributions of the two areas to behaviour through the ratio *a_x_/a_y_*, and (*ii*) one can dissociate how the weight *patterns* (**w**_*x*_ and **w**_*y*_) and their *scales* (*a_x_* and *a_y_*) affect the output of the decoder.

This mathematical separation is also appealing because it provides a common framework to synthesize results from experiments conducted at two fundamentally different levels of granularity. One class of experiments involves making fine measurements such as the correlation between trial-by-trial fluctuations in the activity *r_k_* of an individual neuron *k* and the animal’s decision (**Fig 1a**). The second class of experiments studies causation by measuring behavioural effects of inactivating certain candidate brain areas. For perceptual discrimination tasks, this is done by comparing coarse measures such as the animal’s discrimination thresholds before (*ϑ*) and after (ϑ_*-x*_ and ϑ_-*y*_) inactivating population *x* or *y* (**Figure 1b**).

**Figure 1.**
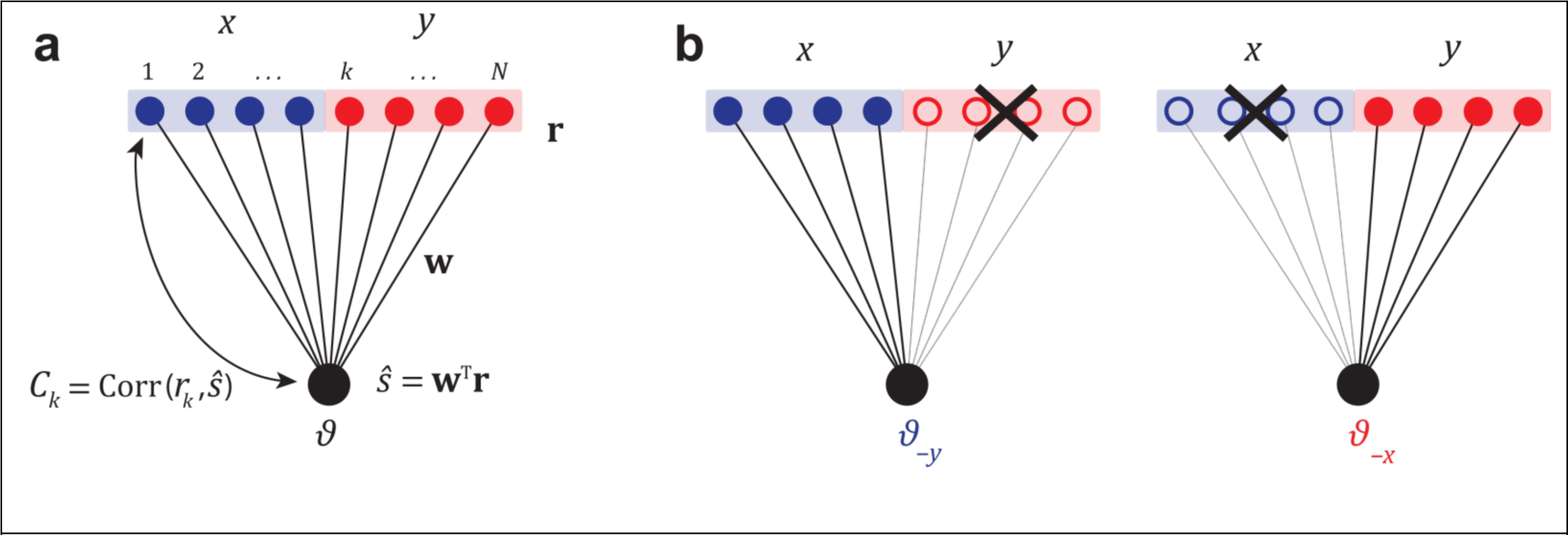
Experimental strategies. **(a)** An illustration of a feedforward network with linear readout. The decoder linearly combines the activity **r** of neurons in populations *x* and *y* with weights **w**, to produce an estimate 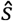 of the stimulus. Activity of individual neurons *r_k_* is correlated with 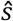 and is quantified by either the choice probability *CP_k_*, or the closely related choice correlation *C_k_*. In an optimal system, the weights **w** generate choice correlations that satisfy **equation 2.1**. **(b)** In inactivation experiments, the neurons from each population are inactivated and the resulting changes in behavioural threshold are recorded.

We would like to use these experimental measurements to identify the relative behavioural contributions of two brain areas. Therefore we will present a technique to infer neuronal weights in two brain areas, focusing primarily on how to extract the scaling factors, *a_x_* and *a_y_*, of the brain areas rather than the fine structure, **w**_*x*_ and **w**_*y*_, of the decoding weights. We first present some results that allow us to examine the pattern of choice correlations of neurons in both areas to characterize the degree of suboptimality in decoding. We will then show how to combine choice correlations with inactivation results to obtain quantitative estimates of the relative scaling of readout weights in those areas.

### Analysis of choice correlations

Choice correlation of a neuron *k* is the correlation coefficient, across repeated trials with the same stimulus *s*, between its response *r_k_* and the animal’s estimate of the stimulus 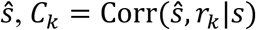. It has recently been shown that readout weights are optimal only if neuronal choice correlations all satisfy the following relation[15] (**Supplementary note S1**):

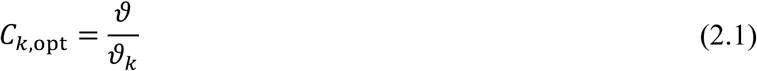

where *C_k,opt_* is the choice correlation of neuron *k* expected from optimal decoding, **ϑ*_k_* is the discrimination threshold of neuron *k*, and *ϑ* is the behavioural discrimination threshold. Therefore if neurons from both areas satisfy the above equation, this gives us strong evidence that the neuronal weights and consequently their relative scales ***a***=(*a_x_*, *a_y_*) are optimal. As we will see later, the exact values of ***a*** can then be directly extracted from the behavioural thresholds *ϑ*_−*x*_ and *ϑ*_−*y*_ following inactivation of those areas.

The pattern of choice correlations generated by any generic suboptimal decoder is more complicated, as it depends explicitly on the structure of noise covariance[14]. For a population of *N* neurons, the covariance *Σ* describes the noise power along *N* orthogonal noise modes. Each of these modes contributes to the overall choice correlation according to (**Supplementary note S2**):

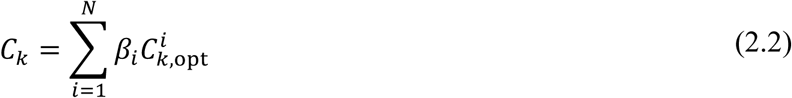

In this expression we have decomposed the optimal pattern of choice correlations *C_k,opt_* into components 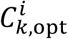 originating from the different noise modes of *Σ*, with 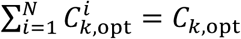. The multipliers *β_i_* reflect the extent of suboptimality. When decoding weights are optimal, every multiplier *β_i_* = 1, so the above equation reduces to **equation 2.1**.

In principle, it is very difficult to estimate all of the multipliers *β_i_* because the components 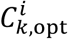 depend on the individual noise modes of *Σ* (**Methods M1 – equation 4**). Directly measuring *Σ* is a notoriously challenging task[20] that involves simultaneously recording the activity of a large population of neurons, and is nearly impossible for certain areas due to the geometry of the brain. Even if such recordings are carried out, it would be impossible to get an accurate assessment of the fine structure of covariance with limited data due to errors arising from finite measurement density[24]. Fortunately, since neuronal choice correlations are measurably large, it follows that one can infer decoding weights with reasonable precision by estimating the few leading multipliers that depend only on the most dominant modes of covariance. This is because if the correlated noise modes with small variance were to dominate the decoder, then only a tiny fraction of each neuron’s variations would propagate to the decision, leading to immeasurably small choice correlations[15] (**Figure S1**). It is possible to determine properties of the leading modes of covariance without large-scale recordings, and we will consider two ways producing two different noise models: *extensive information* and *limited information*.

#### Extensive information model

A common way to measure important components of the covariance structure is through pairwise recordings. Noise covariance measured between pairs of neurons can be modeled as a function of their response properties, such as the difference in their preferred stimulus or the similarity of their tuning functions, to obtain empirical models of noise. One such model is limited-range noise correlations[25–30], so called because they are proportional to signal correlation and thereby limited in range to pairs with similar tuning. We use this model to approximate a full noise covariance for all neurons in the population[31,32] (**Methods M8 — equation 7.1**). Although the resulting covariance matrix is unlikely to capture fine details accurately, if the model is reasonable then most of the variance would be captured by the leading modes.

When decoding two populations *x* and *y*, one has to consider at least two leading modes to capture the two underlying degrees of freedom decoded by scaling factors *a_x_* and *a_y_*. In this minimal case, choice correlations are given by 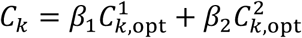. We can compute 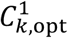 and 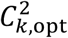 from the leading modes of covariance (**Methods M1** – **equation 4**), and use them to estimate *β*_1_ and *β*_2_ by linear regression. If there are two dominant noise modes and they affect both populations, then we can approximate *Σ* with a rank-two noise covariance matrix composed of both independent (*ϵ*_*xx*_ and *ε*_*yy*_) and correlated (*ε*_*xy*_) noise between the two areas (**Supplementary note S3**). If the two modes were actually uncorrelated, with *ε*_*xy*_=0, so that each mode affects just one population, then the multipliers *β*_1_ and *β*_2_ would be specific to neurons in each population and therefore correspond to *β_x_* and *β_y_*.

A characteristic feature of extensive information models is that the amount of information in the neural activity is very large because it grows with population size[33–35], hence the name. The amount of information extracted by a decoder restricted to the subspace spanned by the few dominant components of covariance cannot be greater than the information available in that subspace. For a model with extensive information, this subset would be a tiny fraction of the total information available in the population. Although this restriction is justified by the large magnitude of neuronal choice correlations, the choice of this noise model is only justified under the assumption that the brain is radically suboptimal.

#### Limited information model

Extensive information models are based on measurements of neural populations but, as we mentioned above, current recordings are not sufficient to measure or even infer the covariance matrix *in vivo*. It is therefore possible that information in cortex is not extensive. Indeed, the extensive information model conflicts with the fact that cortical neurons receive their inputs from a smaller population of neurons. The cortex must then inherit not only the input signal but also any noise in that input. This generates information-limiting correlations[15,20] in cortex, a form of correlated noise that looks exactly like the decoding weights from choice-related activity depends on the noise covariance, we also consider the consequences of information-limiting correlations.

For fine discrimination between two neighboring stimuli *s* and *s*+*δ s*, the signal is given by the change in mean population responses **f**(s+δs)−**f**(s)′δ*s* **f**^′^(*s*). Information-limiting correlations for this task thus fluctuate along the direction **f**′, generating a covariance containing differential correlations[20] — that is, a covariance component proportional to **f**′**f**′^T^. The constant of proportionality, which we denote as *ε*, represents the variance of information-limiting correlations. With increasing population size, both the signal and this noise component grow identically, resulting in no further improvement in signal-to-noise ratio, and thus no improvement in discriminability. In general, *ε* could be very small, and hence information-limiting correlations may be very hard to detect with limited data as they are easily swamped by noise arising from other sources. Nevertheless, this noise has enormous implications for decoding large populations because it limits the total information to 1/*ε*.

When dealing with two populations *x* and *y*, one has to keep in mind that although they may together receive limited information, they need not inherit it from exactly the same upstream neurons. Therefore we construct a more general model allowing the two populations to receive both distinct and shared information. The covariance between two neurons in this more general model would still be proportional to the product of the derivative of their tuning curves. However the constant of proportionality varies depending on whether the pair of neurons are both from the same population *x* (*ε*_*xx*_), both from *y* (*ε*_*yy*_), or from different populations (*ε*_*xy*_) (**Methods M9 – equation 8**). For a large population with this noise structure, the total information content within the *x* and *y* subpopulations alone are by construction equal to 1/*ε*_*xx*_ and 1/*ε*_*yy*_ respectively. The information in both populations together is limited as well, once again by the **f**′**f**′^T^ component of the covariance. Depending on *ε*_*xy*_, the two subpopulations may contain completely redundant, independent, or synergistic information[36,37]. In case the two populations receive information from the same source, then *ε*_*xx*_=*ε*_*yy*_=*ε*_*xy*_ yielding the familiar form of information-limiting correlations[15,20] 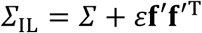. Correlations that limit information within a single neural population introduce redundancy. As a consequence, many different decoding weights can extract essentially the same information. The system is then robust to some suboptimal decoding, which makes it easier to achieve near-optimal behavioural performance[15]. In the noise model for two populations described above, this is also true for each population individually. We can generalize this robustness in our framework by considering separate decoders of each population that produce estimates, 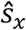 and 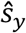, that are near-optimal for their corresponding areas. Importantly, however, these estimates may have different variances, and may even covary, so they need to be properly combined to produce a good single estimate according to **equation 1**. While information-limiting correlations within each area would make the system generally robust to the choice of weight patterns **w**_*x*_ or **w**_*y*_, suboptimality could yet arise from an incorrect scaling (*a_x_* and *a_y_*) of the individual near-optimal estimates. This is because after the dimensionality reduction from large redundant populations down to single unbiased estimates per population, there is no redundancy left: just one degree of freedom remains for the decoder, so different ways of combining the estimates are not equivalent.

If the brain indeed combines activity from different areas suboptimally in this manner, then simplifying **equation 2.2** in the presence of information-limiting correlations gives choice correlations within each area that are not equal to the optimal choice correlations, but are proportional to them (**Supplementary note S5**):

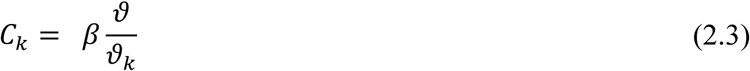

Under these conditions, choice correlations in different areas *x* and *y* may have different multipliers *β*, say *β_x_* and *β_y_*, which depend on the scaling of the two brain areas and on the covariance between the two estimates derived from them. These multipliers *β_x_* and *β_y_* can be directly identified by regressing measured choice correlations against *ϑ*/**ϑ*_k_*, the choice correlations predicted for optimal decoding.

### Combining choice correlations and inactivation effects to infer decoding weights

In the previous section, we showed how to reduce the fine structure of choice correlations down to one number for each population — *β_x_* and *β_y_*. We will now show how these multipliers can be used, together with the behavioural thresholds *ϑ*_−*x*_ and *ϑ*_−*y*_ following inactivation of areas *x* and *y*, respectively, to infer the relative scaling of their weights *a_x_* and *a_y_*. Inactivating an area is equivalent to setting the scaling of weights in that area to zero, so from **equation 1**, the animal’s total estimate 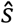 would be equal to either 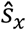 or 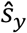, depending on which area is inactivated. The resultant behavioural threshold would simply reflect the variance of the remaining estimate, which is equal to the magnitude of dominant decoded noise within the active area, so 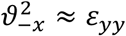 and 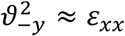. If populations *x* and *y* are uncorrelated (*ε*_*xy*_ = 0), then the ratio of weight scalings can be factorized into a product of ratios (**Supplementary note S6**):

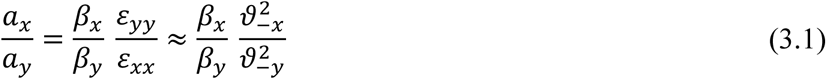

where the two independent factors represent outcomes of correlational and causal studies. If readout is optimal, then the multipliers *β_x_* and *β_y_* are both equal to one, so 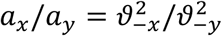. This is consistent with the general belief that the behavioural effects of inactivating a brain area must be commensurate with its contribution to the behaviour. A departure from optimality could break this relationship, so the effects of causal manipulation may not match the relative roles of the brain areas (**Figure S2**). Even in purely feedforward networks, the magnitude of neuronal choice correlations need not equal the effects of inactivation. Thus, disagreements between the two experimental outcomes should not be entirely surprising and do not undermine the functional significance of either.

In fact, **equation 3.1** revealed how one can combine choice correlations and behavioural thresholds to infer the contributions of two uncorrelated areas. But if the areas are correlated, one must explicitly account for the magnitude of correlation between areas *ε*_*xy*_ and the ratio of scales no longer factorizes:

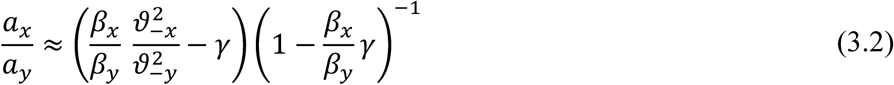

where *γ* = *ε*_*xy*_/*ε*_*xx*_ is the magnitude of correlated noise between the two populations’ estimates relative to the variance of estimates from *x* alone. Note that one can also use **equations 3.1** and **3.2** to compute the optimal weight scaling factors simply by setting both *β_x_* and *β_y_* to 1. Therefore we can use these equations not only to determine the relative weights of brain areas but to also to evaluate precisely how suboptimal those weights are.

### Application to data

We now use the techniques developed so far to infer the relative contributions of two brain areas in macaque monkeys to heading discrimination. Data were collected from monkeys trained to discriminate their direction of self-motion in the horizontal plane (**Figure 2a**) using vestibular (inertial motion) and/or visual (optic flow) cues (**Methods M4**; see also refs. [21,23]). At the end of each trial, the animal reported whether their perceived heading 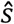 was leftward (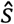 < 0°) or rightward (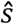 > 0°) relative to straight ahead.

**Figure 2.**
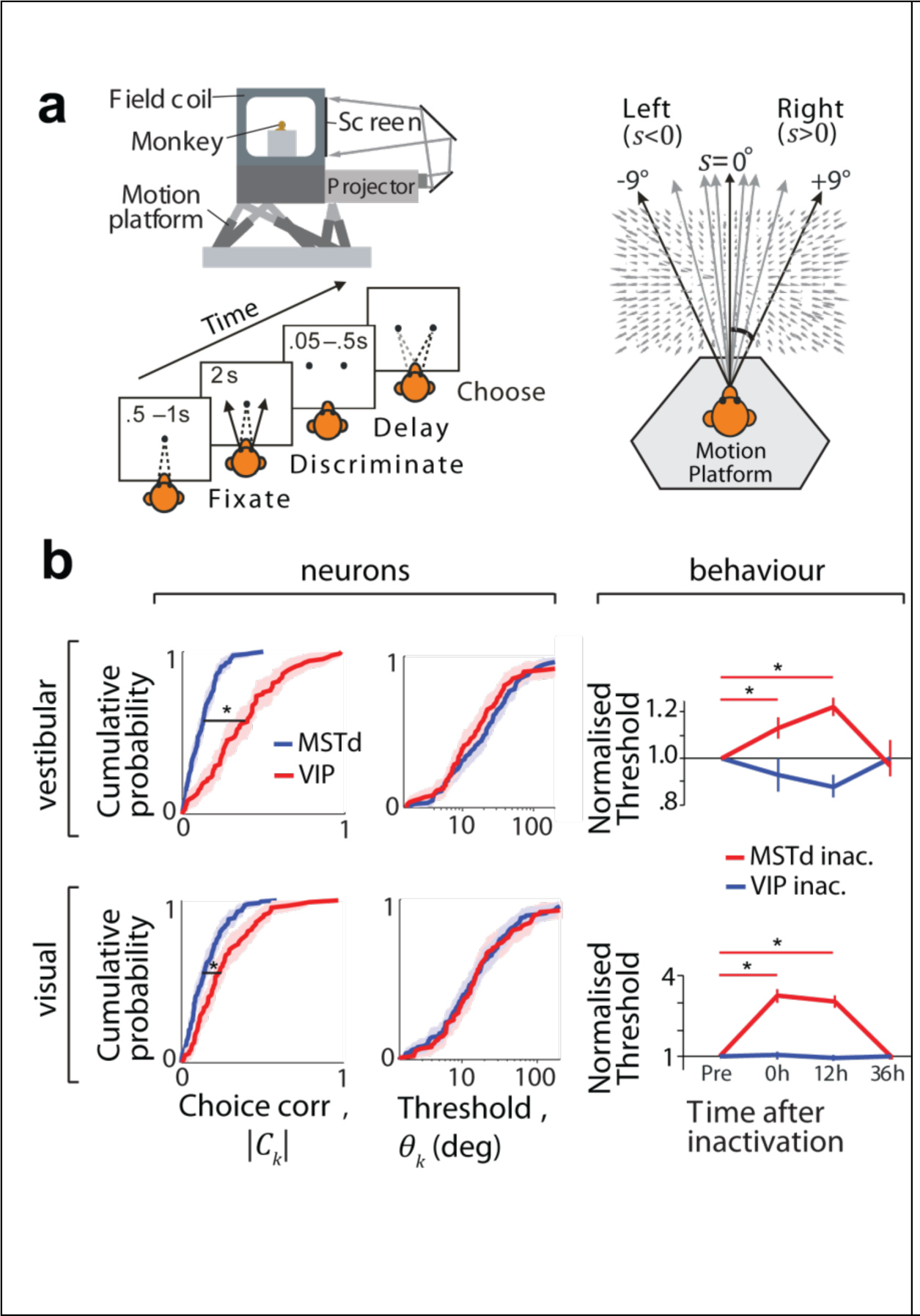
Choice-related activity and effects of inactivation. **(a)** Behavioural task: the monkey sits on a motion platform facing a screen. He fixates on a small target at the center of the screen, and then we induce a self-motion percept by moving the platform (vestibular condition) or by displaying an optic flow pattern on the screen (visual condition). The fixation target then disappears and the monkey reports his percept by making a saccade to one of two choice targets. **(b) Left**: Neurons in both MSTd (*n* = 129) and VIP (*n* = 88) exhibited significant choice correlations (CCs). The median CC of VIP neurons was significantly greater than that of MSTd neurons (*;*p*<0.001, Wilcoxon rank-sum test) in both vestibular (top) and visual (bottom) conditions. **Middle**: Median neuronal thresholds were not significantly different between areas (vestibular: *p* = 0.94, visual: *p* = 0.86, Wilcoxon rank–sum test). **Right**: Average discrimination thresholds at different times relative to inactivation of VIP and MSTd. All threshold values were normalized by the corresponding baseline thresholds (“pre”). Shaded regions and error bars denote standard errors of the mean (SEM); asterisks indicate significant differences (\**p*<.05, *t*–test). Neural data re-analyzed from refs. [21,23]. Inactivation data reproduced from refs. [18,22].

### Discrepancy between correlation and causal studies

Responses of single neurons were recorded from either area MSTd (monkeys A and C; *n* = 129) or area VIP (monkeys C and U; *n* = 88) during the heading discrimination task (**Methods M5**). Basic aspects of these responses were analyzed and reported in earlier work[21,23]. Briefly, it was found that neurons in VIP had substantially greater choice correlations (CC) than those in MSTd (**Figure 2b –** left) for both the vestibular and visual conditions. This difference in CC between areas could not be attributed to differences in neuronal thresholds *ϑ_k_* (**Figure 2b –** middle), defined as the stimulus magnitude that can be discriminated correctly 68% of the time (*d*′ = 1) from neuron *k*’s response *r_k_* (**Methods M6**; **Figure S3**). Based on its greater CCs, one might expect that VIP plays a more important role in heading discrimination than MSTd. In striking contrast to this expectation, a recent study showed that there was no significant change in heading thresholds following VIP inactivation for either the visual or vestibular stimulus conditions[18] (**Figure 2b** – right (blue); monkeys B and J). On the other hand, inactivation of MSTd using a nearly identical experimental protocol led to substantial deficits in heading discrimination performance[22] (**Figure 2b** – right (red); monkeys C, J, and S). The neural and inactivation studies in VIP used non-overlapping subject pools, so the observed dissociation between CCs and inactivation effects could potentially reflect the idiosyncrasies of the subjects’ brains. To rule this out, we repeated the inactivation experiment by specifically targeting Muscimol injections to sites in area VIP that were previously found to contain neurons with high CCs in another monkey and obtained similar results (**Figure S4**).

These findings reveal a striking dissociation between choice correlations and effects of causal manipulation: VIP has much greater CCs than MSTd yet inactivating VIP does not impair performance. One may be tempted to simply conclude that VIP does not contribute to heading perception. We will now show that this is not necessarily true. Depending on the structure of correlated noise and the decoding strategy, neurons in both areas may be read out in a manner that is entirely consistent with the observed effects of inactivation.

## Test for Optimality

We first asked if the above results can simply be explained if the brain allocated weights optimally to the two areas. To answer this, we tested if neuronal choice correlations satisfied **equation 2.1**. Binary discrimination experiments typically do not measure choice correlations 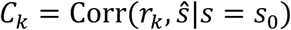 because they do not have direct access to the animal’s continuous stimulus estimate 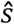; they only track the animal’s binary choice. Instead they measure a related quantity known as choice probability defined as the probability that a rightward choice is associated with an increase in response of neuron *k* according to 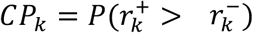 where 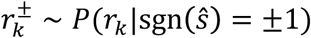 is a response 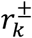 of neuron *k* when the animal chooses ±1. Therefore we first transformed the measured choice probabilities to choice correlations using a known relation[14] before further analyses (**Methods M7**). Equivalently, one could measure the correlation 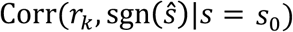 between the neural response and the binary choice, which^15^ showed is ≈0.8 *C_k_*. Note that the above definition gives choice correlations that are either positive or negative depending on whether a rightward choice is associated with an increase or decrease in neuronal response. Therefore we adjusted **equation 2.1** to generate predictions for optimal CCs that accounted for our convention (**Methods M7**).

**Figure 3** compares experimentally measured CCs against the CCs predicted by optimal decoding for all neurons recorded in the vestibular (left panel) and visual (right panel) conditions. Our data are consistent with optimal decoding of MSTd, since the predicted and measured CCs are significantly correlated (vestibular: Pearson’s *r* = 0.65, *p* < 10^−3^; visual: *r* = 0.70, *p* < 10^−3^) with a slope not significantly different from 1 (vestibular: slope = 1.11, 95% confidence interval (CI) = [0.83 1.54]; visual: slope = 1.24, 95% CI = [0.94 1.78]). For VIP, although the predicted and measured CCs are again strongly correlated (vestibular: *r* = 0.80, *p* < 10^−3^; visual: *r* = 0.75, *p* < 10^−3^), the regression slope deviates substantially from unity (vestibular: slope = 2.37, 95% CI = [1.97 3.08]; visual: slope = 1.98, 95% CI = [1.41 2.74]), demonstrating that our data are inconsistent with optimal decoding. Note that, if VIP is decoded suboptimally, this implies that the overall decoding—one based on both VIP and MSTd—is suboptimal as well because the decoder failed to use all information available in the neurons across both populations.

**Figure 3.**
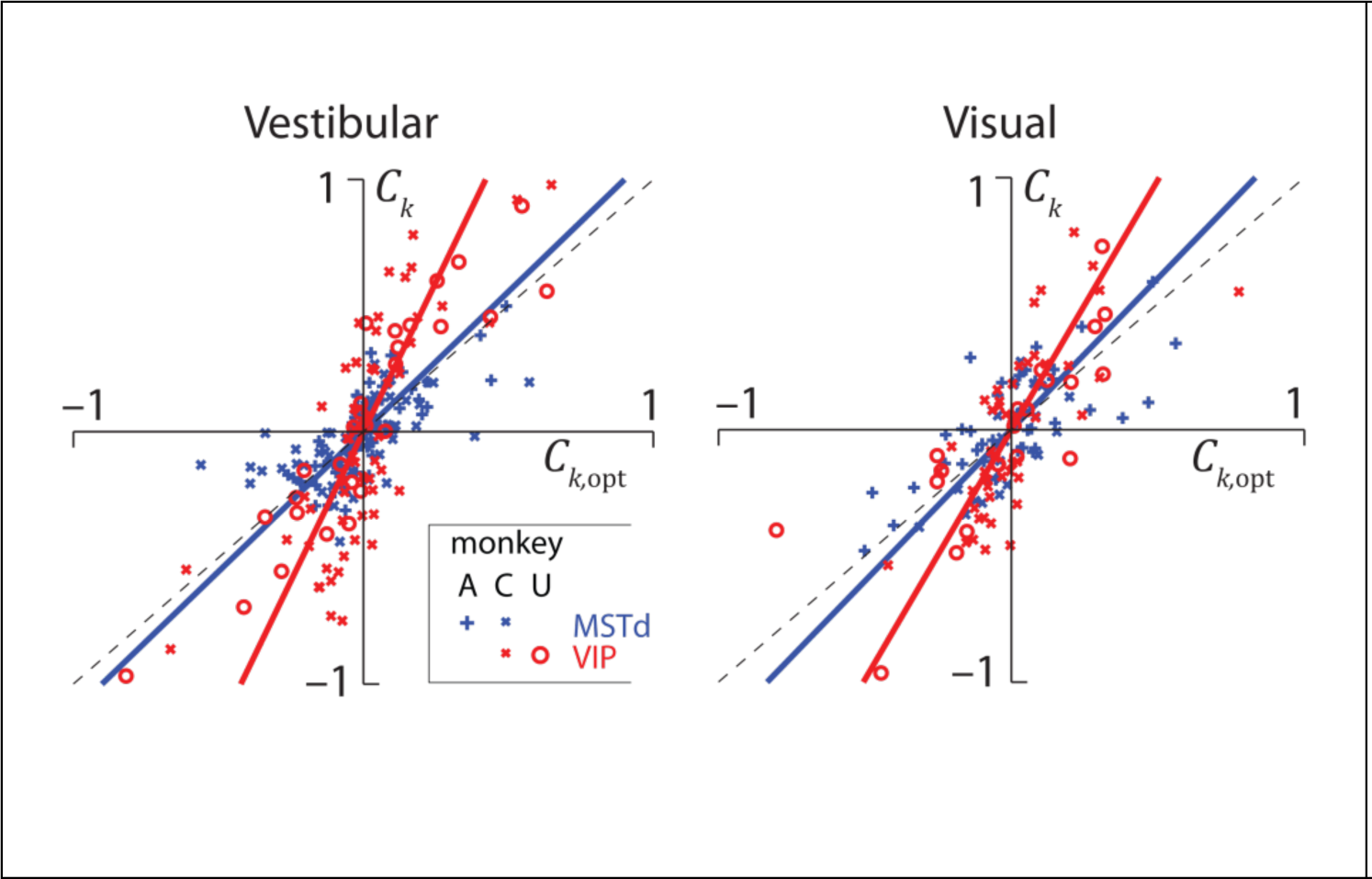
Readout is not optimal. Whereas the experimentally measured choice correlations (*C_k_*) of neurons in MSTd (blue) for both the vestibular (left) and the visual (right) condition are well described by the optimal predictions (*C_k,opt_*), those of VIP neurons are systematically greater (red). This observation was consistent across all monkeys (see **Supplementary Figure S4a** for monkey *X*). Solid lines correspond to the best linear fit. Vestibular data replotted from Ref.[15] with different sign convention (**Methods M7**).

This leads to two questions: First, how much information is lost by suboptimal decoding? Second, how is this information lost? To get precise answers, we will now determine how the brain weights activity in MSTd and VIP to perform heading discrimination.

### Inferring readout weights

Throughout this section, we use subscripts *M* and *V* to denote MSTd and VIP instead of the generic subscripts *x* and *y* used to describe the methods. For clarity, we will restrict our focus to the vestibular condition but results for the visual condition are presented in the supplementary notes. In order to determine decoding weights, we constructed two kinds of covariance structures that implied either extensive or limited information as explained earlier.

In the extensive information case, we modeled noise covariance using data from pairwise recordings within MSTd and VIP reported previously [21,29]. Those experiments established that noise correlation between neurons in these areas tends to increase linearly with the similarity of their tuning functions, or signal correlation (**Methods M8 – equation 7.1**). This relationship between noise and signal correlations has a substantially steeper slope in VIP than in MSTd (MSTd: *m_M_* = 0.19±0.08; VIP: *m_V_* = 0.70±0.16, **Figure S5**). We used these empirical relationships to extrapolate noise correlations between all pairs of independently recorded neurons within each of the two populations, using only their tuning curves, and assuming that any stimulus-dependent changes in correlation were negligible. Since correlations between VIP and MSTd populations were not measured experimentally, we explored different correlation matrices (**Methods M8 – equation 7.2**).

In the limited information case, we added correlations that limited the total information content across the two populations (**Methods M9** – **Equation 8**). For this latter case, we relied on behavioural thresholds before and after inactivation, and choice correlations, to determine the magnitudes of noise within (*ε_MM_* and *ε_VV_*) and between (*ε_MV_*) areas (**Methods M9**). In both cases, we constructed covariances for many different population sizes *N* by sampling equal numbers of neurons from both areas with replacement. The choice of distributing neurons equally among the two areas was made only for convenience and has no bearing on the result as explained later.

**Figure 4a** shows example covariance matrices for both extensive and limited information models for a population of 128 neurons. The two structures look visually similar because the additional fluctuations caused by information-limiting correlations are quite subtle. Nevertheless, there is a huge difference between the two models in terms of their information content (**Figure 4b**). The extensive model has information that grows linearly with *N*, implying that these brain areas have enough information to support behavioural thresholds that are orders of magnitude better than what is typically observed. However when information-limiting correlations are added, information saturates rapidly suggesting that behavioural thresholds may not be much lower than population thresholds even if the decoding weights are fine-tuned for best performance. We will now infer scaling factors *a_M_* and *a_V_* of decoding weights using both noise models and examine their implications.

**Figure 4.**
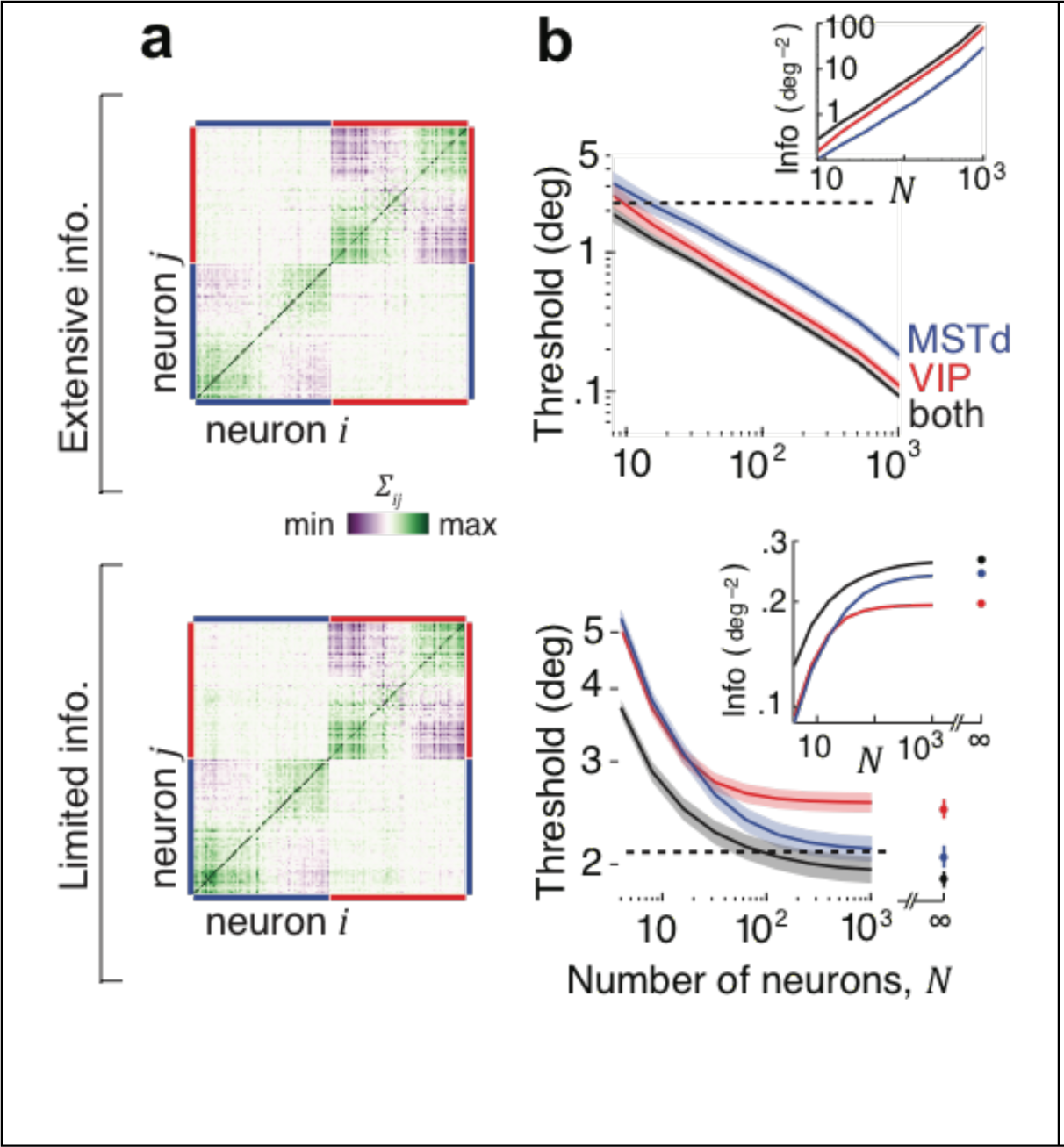
Covariance structure of Extensive and Limited information models. (**a**) Matrix of covariances *Σ*_*ij*_ among neurons in MSTd and VIP (*N* = 128). Top: Extensive information model constructed by sampling according to the empirical relationship in **Supplementary Figure S5**, for the case when the two areas are uncorrelated on average. Bottom: Limited information model adds a small amount of information-limiting correlations with magnitudes (*ε_MM_* = 4.2, *ε_VV_* = 7, *ε_MV_* = 0) chosen arbitrarily for illustration. **(b)** Inset shows the effect of population size on the information content implied by the two kinds of noise in MSTd (blue), VIP (red) and in both areas together (black). If decoded optimally, behavioural thresholds implied by the extensive information model would decrease with *N* resulting in performance levels that are vastly superior to those actually observed in monkeys (black dashed line). Information-limiting correlations cause information to saturate with *N*.

#### Extensive information model

We’ve already seen that the pattern of choice correlations is not consistent with optimal decoding of MSTd and VIP. In fact for the extensive information model, optimal decoding will lead to extremely small CCs by suppressing response components that lie along the leading noise modes as they have very little information (**Figure S6a**). Ironically, the magnitude of CCs found in our data could only have emerged if the response fluctuations along those leading modes substantially influenced animal’s choice (**Figure S6b**). This means that the decoder must be largely confined to the subspace spanned by those modes. We therefore restricted our focus to the two leading eigenvectors **u**^1^ and **u**^2^ of the covariance matrix. When the two populations are uncorrelated, these vectors lie exclusively within the one-dimensional subspaces spanned by neurons in MSTd and VIP respectively (**Figure 5a**). In our case, vectors **u**^1^ and **u**^2^ corresponded to **u**^*V*^ and **u**^*M*^.

**Figure 5.**
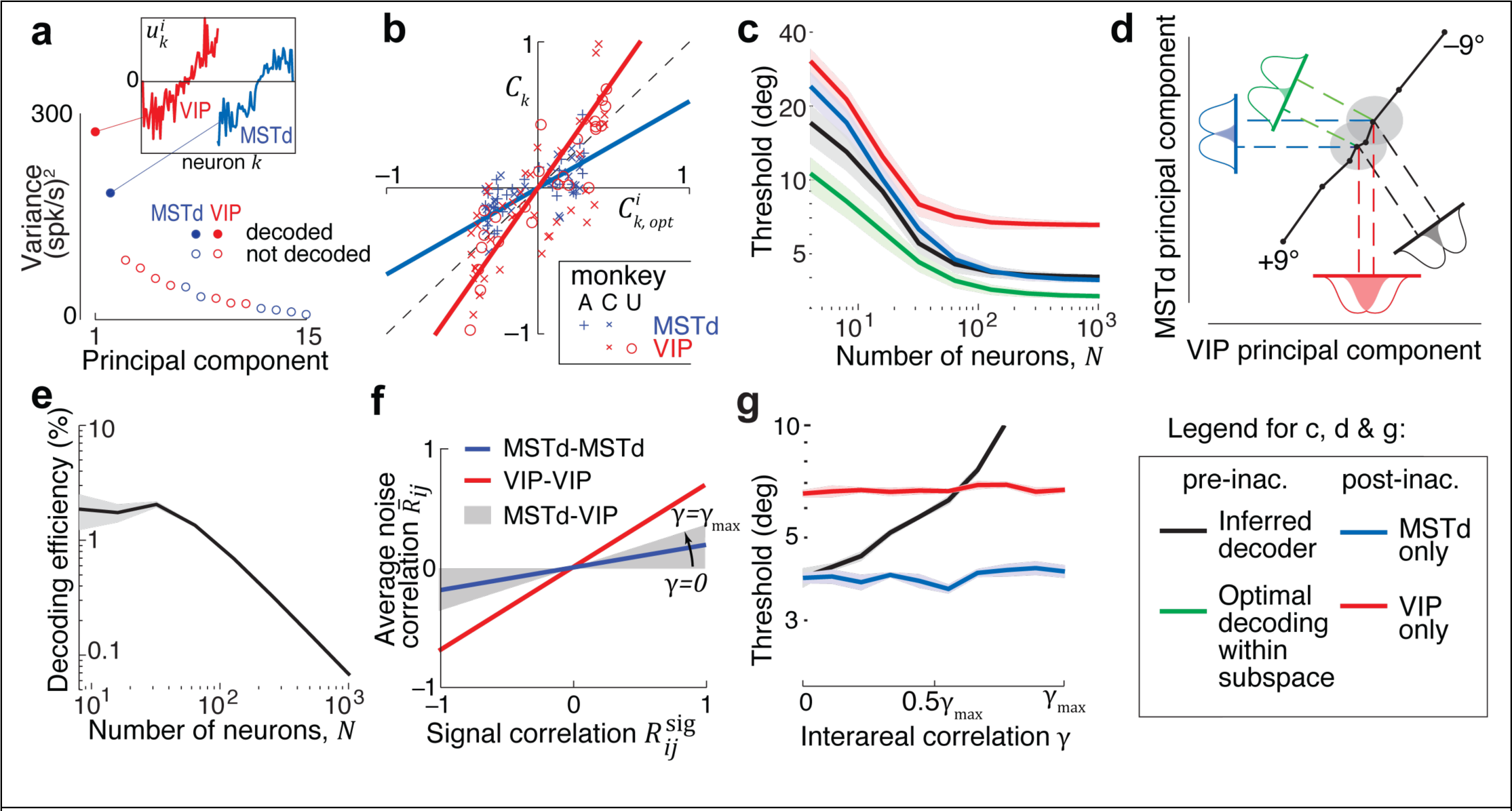
Decoder inferred using the extensive information model. (**a**) Decoding weights were inferred in the subspace of 2 leading principal components of noise covariance (solid circles). Inset: These components lie entirely within the space spanned by neurons in one of the two brain regions. Components are color coded according to the brain region that it inhabits (red=VIP; blue=MSTd). (**b**) Experimentally measured choice correlations (*C_k_*) of individual neurons in VIP (red) and MSTd (blue) are plotted against their respective components 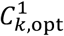 and 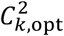 of choice correlations generated from optimally decoding responses within the subspace of 2 leading principal components. (**c**) Unlike the optimal decoder in **Figure 4b**, the behavioural threshold predicted by the inferred weights (black) saturates at a population size of about 100 neurons. The green line indicates the performance of an optimal decoder within the two-dimensional subspace. Inactivating VIP is correctly predicted to have no effect on behavioural performance for large *N* (blue), while MSTd inactivation increases the threshold (red). (**d**) A schematic of the inferred decoding solution projected onto the first principal component of noise in VIP and MSTd. The solid colored lines correspond to the readout directions for the four cases shown in (c). The long diagonal black line is the projection of the mean population responses for headings from −9° to +9°, and the two gray ellipses correspond to the noise distribution at heading directions of ±2°. The colored gaussians correspond to the projections of this signal and noise onto each of the four readout directions, and the overlap between these gaussians corresponds to the probability of discrimination errors. (**e**) The percentage of available information read out by the inferred decoder (the decoding efficiency) decreases with population size, because the decoded information saturates while the total information is extensive. (**f**) Correlations between MSTd and VIP were not measured experimentally. We modeled these correlations according to the same linear trend that on average described correlations within each population, but with different slopes, yielding different interareal correlations parametrized by *γ* = *ϵ_MV_/ϵ_MM_* (**Methods M8 – Equation 7.2**). This slope reaches its maximum allowable value 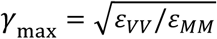, the geometric mean of the slopes for MSTd and VIP. (**g**) For each value of γ, we used the resultant covariance and CCs to infer the decoder, and plotted its behavioural thresholds. Thresholds are shown for a population of 256 neurons, by which point the performance had saturated to its asymptotic value for all γ. Shaded regions in (c), (e), and (g) represent ±1 SEM.

Although decoding only this subspace is not optimal with respect to the total information content in the two areas, a decoder could still be optimal within that subspace. To test this, we estimated the choice correlations 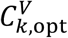 and 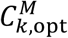 that would be expected from optimally weighting the two areas within this subspace (**Methods M1 – equation 4**). The observed CCs were proportional (MSTd: Pearson’s *r* =0.55, *p* < 10^−3^; VIP: *r* = 0.76, *p* < 10^−3^) to these optimal predictions implying that the leading noise modes of the extensive information model are able to capture the basic structure of choice-related activity in both areas (**Figure 5b**). However the slopes *β_M_* and *β_V_* were significantly different from 1 (*β_M_* = 0.73, 95% CI = [0.63 0.84]; *β_V_* = 2.38, 95% CI = [2.2 2.57]) implying that the weight scalings *a_M_* and *a_V_* must be suboptimal even within the two-dimensional subspace. Since we knew the magnitudes of *ε_MM_* and *ε_VV_* for this noise model from pairwise recordings (**Table 1**), we applied the exact rather than approximate form of **equation 3.1** and obtained a scaling ratio *a_M_*/*a_V_* = 0.8±0.1.

**Table 1.**
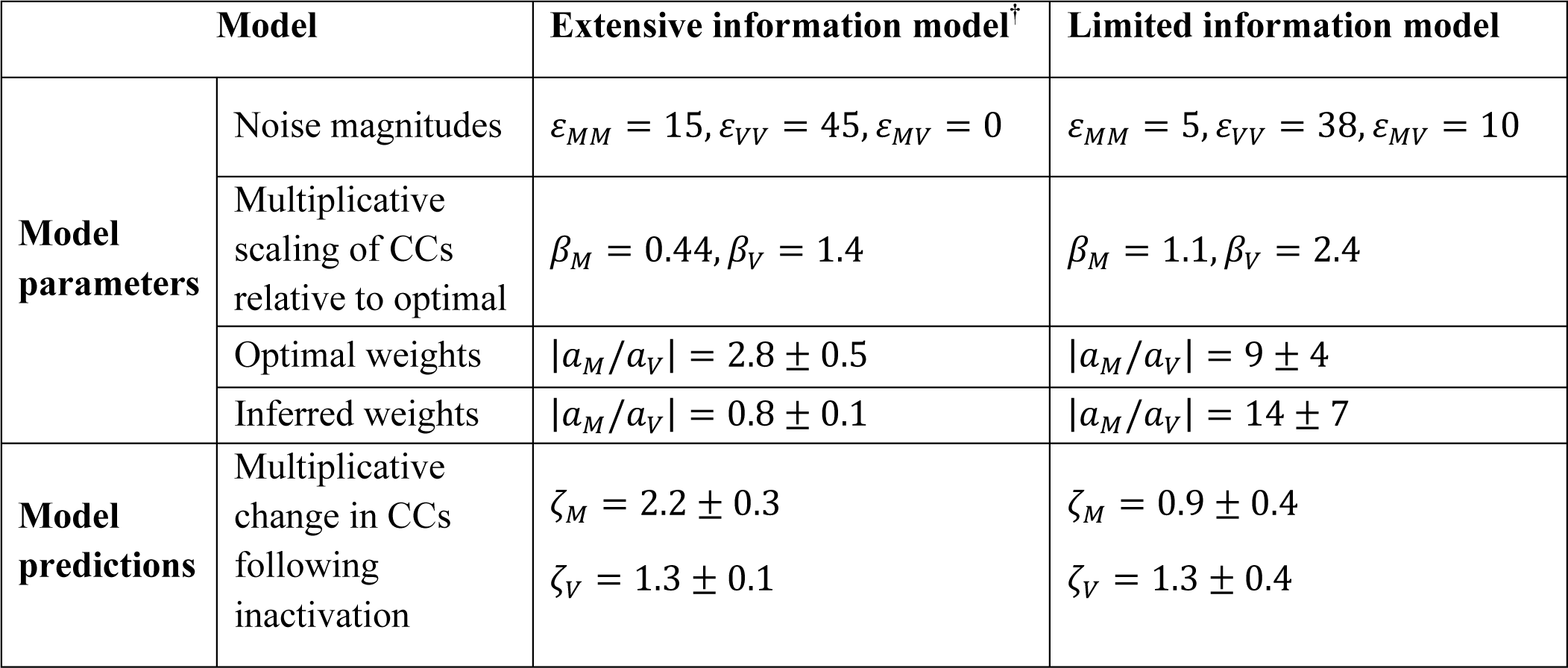
Model parameters and predicted changes in CCs following inactivation for the two covariance models, shown as median±central quartile range. (**^†^**Values correspond to when decoder is inferred using a rank-two approximation of the covariance.)

To test whether the inferred scaling was meaningful, we compared behavioural thresholds implied by the resulting decoding scheme against experimental findings of inactivation. The threshold prior to inactivation is related to the variance of the estimator whose decoding weights **w** are along the direction specified by *a_M_***u**^*M*^ + *a_V_***u**^*V*^. Inactivating either area is equivalent to setting the corresponding scaling to zero so post-inactivation thresholds are given by the variance along the leading noise mode specific to the active area (**u**^*M*^ or **u**^*V*^). We computed pre and post-inactivation thresholds and found they were qualitatively consistent with experimental results: for large populations, MSTd inactivation is predicted to produce a large increase in threshold (**Figure 5c**, red vs black) whereas VIP inactivation is predicted to have little or no effect (**Figure 5c**, blue vs black; see **Figure S7** for visual condition). This correspondence to experimental inactivation results is remarkable because the procedure to deduce scalings *a_M_* and *a_V_* was not constrained in any way by behavioural data, but rather informed entirely by neuronal measurements. We also confirmed that the threshold expected from optimal scalings (**Table 1**) was smaller than that produced by inferred weights (**Figure 5c**, green vs black) implying that the brain indeed weighted the two areas suboptimally.

The above findings are explained graphically in **Figure 5d** by projecting the relevant quantities (tuning curves **f**(*s*), noise covariance *Σ*, decoding weights **w**) onto the subspace of the first two principal components (**u**^*M*^ and **u**^*V*^) of the noise covariance *Σ*. The colored lines indicate different readout directions, determined by the scaling (*a_M_* and *a_V_*) of weights for the two populations. A ratio of ϵ*a_M_*/*a_V_*ϵ > 1 corresponds to greater weight on the estimate derived from MSTd activity, and the associated readout direction will be closer to the principal component of MSTd. The response distributions are depicted as gray ellipses (isoprobability contours) for the two stimuli to be discriminated. The discrimination threshold for different decoders can be obtained simply by projecting these ellipses onto the readout direction of the specified decoder and examining the overlap between the projections. Within this subspace, the ratio ϵ*a_M_*/*a_V_*ϵ of the decoder inferred from CCs was much smaller than the optimal ratio (**Table 1**), meaning that MSTd was given too little weight. Consequently, the response distributions have more overlap along the direction corresponding to the decoder inferred from neuronal CCs (black) than along the optimal direction in that subspace (green). This means that the outputs are less discriminable and thus that the decoding is suboptimal. VIP inactivation (*a_V_* = 0) corresponds to decoding only from MSTd (blue). This happens to produce no deficit because the overlap of the response distributions is similar to that along the original decoder direction. On the other hand, inactivating MSTd (*a_M_* = 0) corresponds to decoding only from VIP (red), where the two response distributions have greater overlap leading to a larger threshold.

It is important to keep in mind that decoding the noisiest two-dimensional subspace, which throws away all signal components in the remaining low-noise *N*–2 response dimensions, is a much more severe suboptimality than misweighting the two areas’ signals within that restricted subspace, which loses less than half the information (**Figure 5c**). As illustrated in **Figure 5e**, the fraction of available information recovered by this decoder (*η*) drops precipitously with the number of neurons (*η* ~ 2.5 *N*^−1^). Moreover, for this model, a steeper relationship between signal and noise correlations leads to greater CCs. This is because the model is only consistent with suboptimal decoding that fails to remove the strong noise correlations; these noise correlations are decoded to drive the choice, and thus correlate neurons not only with each other but also with that choice. Thus, in the extensive information model, high CCs are a consequence of decoding a restricted subspace of neural activity, a radically suboptimal strategy for the brain.

Behavioural predictions of this model were robust to assumptions about the exact size of the decoded subspace (**Figure S8**), but were found to depend on the magnitude of noise correlations between the VIP and MSTd populations. Since interareal correlations were not measured, we systematically varied the strength of these correlations by changing *γ* (**Figure 5f**), and used **equation 3.2** to infer weight scalings for each case. We used these scalings to generate behavioural predictions for different values of *γ*. Predictions for one example value of these correlations are shown in **Figure S9**. Behavioural predictions progressively worsened as a function of the strength of noise correlations between MSTd and VIP: for this model, even weak but nonzero interareal correlations imply that inactivating area VIP should improve behavioural performance (**Figure 5g**).

#### Limited information model

In the presence of information-limiting correlations, choice correlations must be proportional to the ratio of behavioural to neuronal thresholds (**Equation 2.3**). This was indeed the case both in MSTd and VIP as we showed already in **Figure 3**. Those slopes correspond to the multipliers *β_M_* and *β_V_* for this model, and were found to be different for the two areas (**Table 1**).

As we noted earlier, unlike the leading modes of noise in the extensive information model, the magnitudes of information-limiting correlations (*ε_MM_*, *ε_VV_*, and *ε_MV_*) are difficult to measure. Nevertheless, we can deduce them from behaviour because behavioural precision is ultimately limited by these correlations. Briefly, using behavioural thresholds *after* inactivation of each area, along with *β_M_* and *β_V_* derived from choice correlations as additional constraints, we can simultaneously infer the magnitude of information-limiting correlation within each area (*ε_MM_* and *ε_VV_*), the correlated component of the noise (*ε_MV_*), and weight scalings (*a_M_* and *a_V_*) (**Methods M9**). A model based on these inferred parameters correctly predicted that the behavioural threshold *before* inactivation would not be significantly different from threshold following VIP inactivation (**Figure 6a**; see **Figure S10** for visual condition). This was because the scaling of weights in MSTd was much larger than in VIP according to this model (*a_M_* » *a_V_*, **Table 1**), so inactivating VIP had little impact on the output of the decoder and left behaviour nearly unaffected. Unlike the decoder inferred for the extensive information model, the efficiency *η* of this decoder did not depend on the size of the population being decoded (**Figure 6b**, vestibular: *η* = 79 ± 13%) because neurons in this model carry a lot of redundant information.

**Figure 6.**
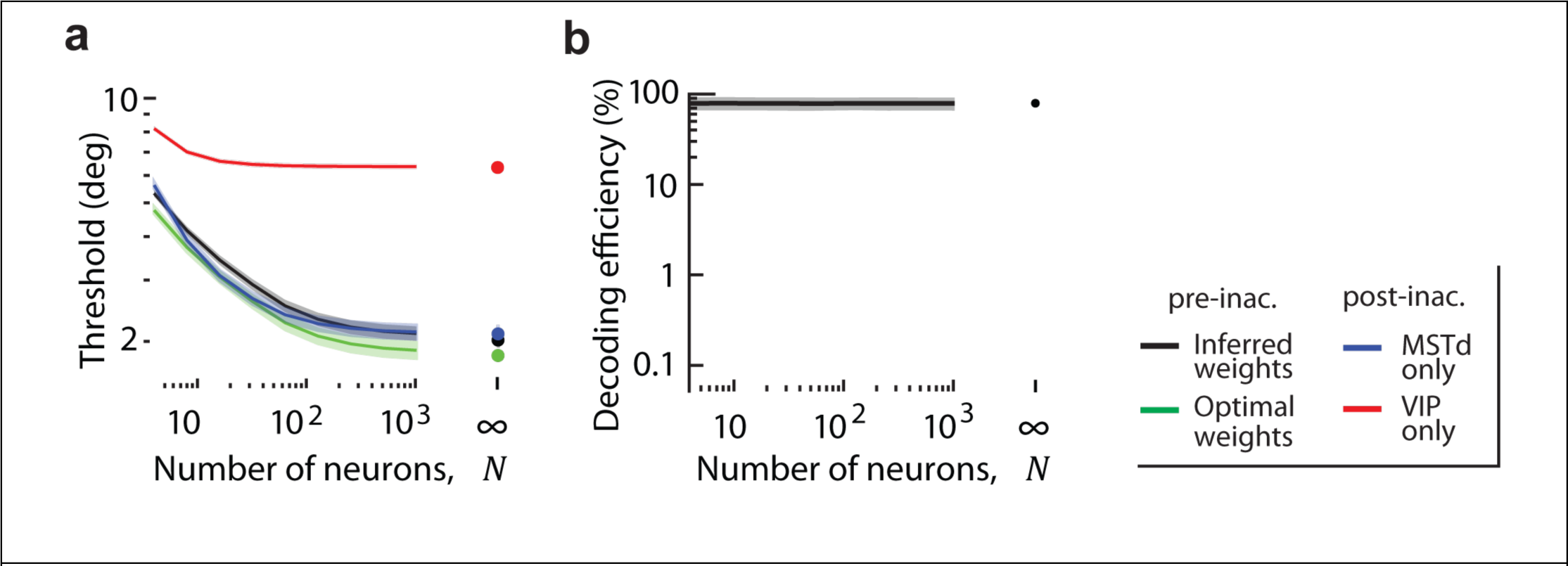
Decoder inferred using the limited information model. (a) Like decoding in the presence of extensive information, this decoder is suboptimal (black vs green), and can account for the behavioural effects of inactivation. **(b)** Unlike decoding in the extensive information model, the efficiency of this decoder is high and insensitive to population size. Shaded areas represent ±1 SEM.

All analyses above were performed on neural data in the central 400ms of the trials following earlier work. However our conclusions are robust to the specific time (**Figure S11**) and duration (**Figure S12**) of the analysis window. Additionally, although we extrapolated our data to larger populations by resampling from a set of about 100 neurons recorded from each area, our results are not attributable to the limited size of the recording (**Figure S13**). We also extended our model to account for the fact that the two brain areas may have only been partially inactivated by Muscimol, and found that our conclusions hold under a wide range of partial inactivations (**Supplementary note S8**; **Figure S14**). Finally, we assumed that inactivation leaves responses in the un-inactivated area unaffected, as would be the case in a purely feedforward network model. While an exhaustive treatment of recurrent networks is beyond the scope of this work, we find that our conclusions can still hold if the above assumption is compromised by recurrent connections between MSTd and VIP (**Supplementary note S9**; **Figure S15**).

#### Comparison of the two decoding strategies

We inferred decoding weights in the presence of two fundamentally different types of noise, the extensive information model and the limited information model. Both of these decoders could account for the behavioural effects of selectively inactivating either MSTd or VIP, albeit with very different readout schemes. For the extensive information model, neurons in area VIP were weighted more heavily than optimal, and vice-versa in the presence of information-limiting noise (**Table 1, Figure 7a**). Why do the two models have such different weightings? Both noise models have larger noise in VIP than MSTd, but differ in correlations between the two areas. In the extensive information model, the interareal correlations must be nearly zero to be consistent with behavioural data (**Figure 5g** and **Figure S9**), and the neuronal weights in VIP must be high to account for the high CCs. In the limited information model, the significant interareal correlations explain the large CCs in VIP, even with a readout mostly confined to MSTd.

**Figure 7.**
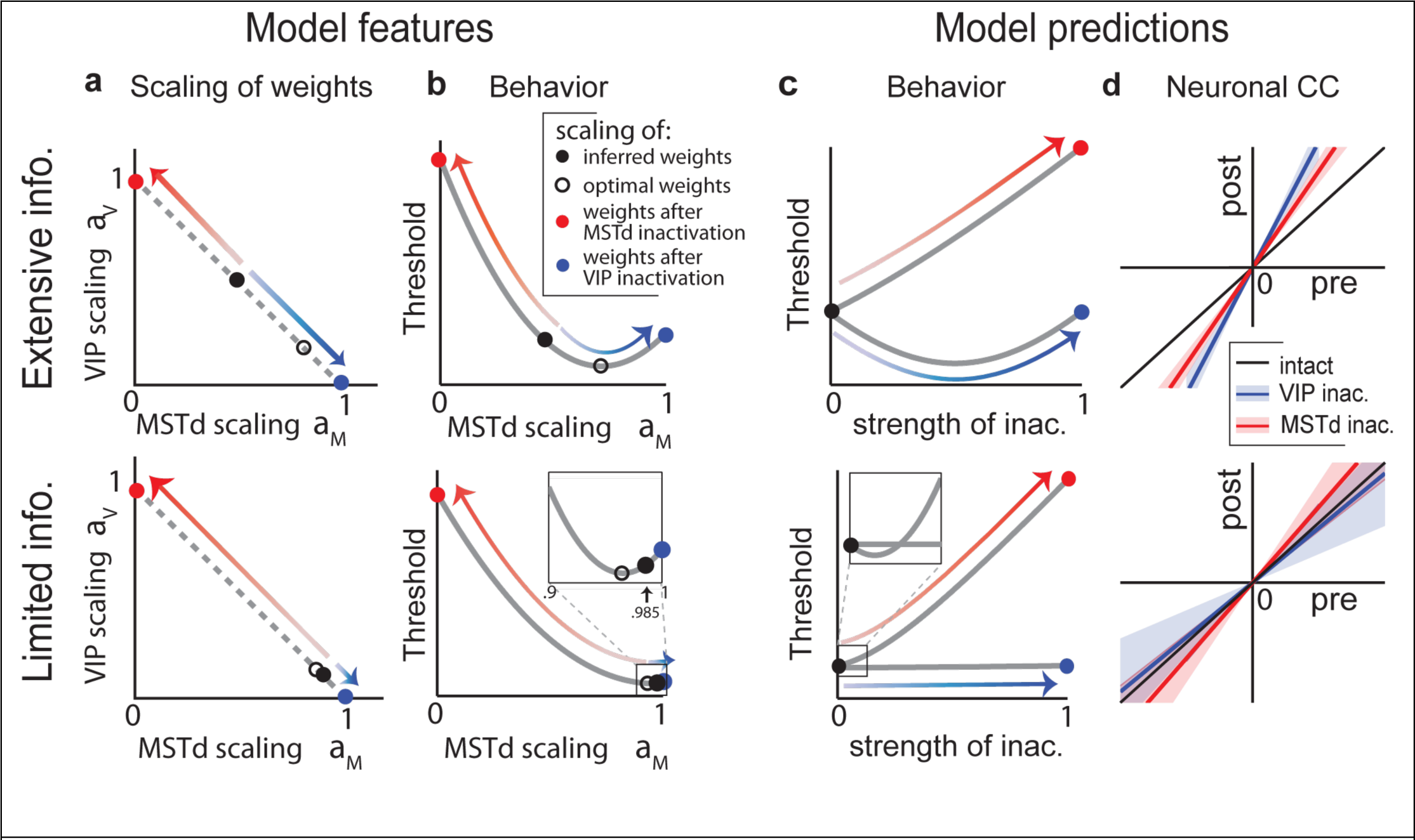
Decoding strategy and model predictions for the extensive information model and the limited information model. (**a**) Optimal (open black) and inferred (filled black) scaling of weights in MSTd (*a_M_*) and VIP (*a_V_*). Inactivation of either MSTd (red) or VIP (blue) confines the readout to the active area resulting in a scaling of 1. Red and blue arrows indicate the transformation resulting from inactivating MSTd and VIP respectively. The scaling factors always sum to 1. **(b)** Behavioural threshold *ϑ* as a function of *a_M_*. Whereas *ϑ* increases following MSTd inactivation for both models (red), it improves initially following partial VIP inactivation (blue) in the extensive information model (top) but remains unchanged in the limited information model (bottom). (**c**) The same curves can be replotted as a function of the strength of inactivation of MSTd (red) or VIP (blue) yielding behavioural predictions for partial inactivation of the areas. **(d)** Choice correlations (CC) of neurons in MSTd (blue) and VIP (red), before and after inactivation of VIP and MSTd respectively. Again the results following MSTd inactivation do not discriminate the two information models, but for VIP inactivation the predictions differ, showing increased CCs for the extensive information model and decreased CCs for the limited information model. Slopes of the lines correspond to ζ*_M_* and ζ_*V*_ in **Equation (9)**, and shaded regions indicate ±1 s.d. of uncertainty.

How could such fundamentally different strategies lead to the same behavioural consequences? For a given noise model, an optimal decoder achieves the lowest possible behavioural threshold by scaling the weights of neurons in the two areas according to a particular optimal ratio *a_M_*/*a_V_*. Ratios that are either smaller or larger than this optimum will both result in an increase in the behavioural threshold due to suboptimality. This produces a *U-shaped* performance curve. Under certain precise conditions, complete inactivation of one of the areas will leave behavioural performance unchanged, exactly on the other side of the optimum. This is the case for VIP according to the extensive information model (**Figure 7b – top**). On the other hand, if the weight is already too small to influence behaviour then inactivation may not appreciably change performance, as demonstrated by the limited information model (**Figure 7b – bottom**).

#### Model predictions

According to the extensive information model, the brain loses almost all of its information by poorly weighting its available signals. Moreover, even beyond this poor overall decoding, the model brain gives VIP too much weight. As a consequence, this model makes a counterintuitive prediction that gradually inactivating VIP should *improve* behavioural performance! A hint of this might already be seen in **Figure 1d** and **Figure S4b** for the vestibular condition (both 0 and 12 h), although the difference was not statistically significant. Beyond a certain level of inactivation, as the weight decreases past the optimal scaling of the two areas, performance should worsen again (**Figure 7c – top**). According to the extensive information model, the brain just so happens to overweight VIP under normal conditions by about the same amount as it underweights VIP after inactivation. Suboptimal decoding in the limited information model has the opposite effect, giving too little weight to VIP, while overweighting MSTd. However, according to this model, the available information in VIP is small, because when MSTd is inactivated the behavioural thresholds are substantially worse (**Figure 7c – bottom**). Thus the suboptimality due to underweighting VIP is mild (around 80% in both visual and vestibular conditions, as described above), and the predicted improvement following partial MSTd inactivation is negligible as gradual inactivation quickly shoots past the optimum. Graded inactivation of brain areas can be accomplished by varying the concentration of muscimol, as well as the number of injections. In fact, we have previously reported that behavioural thresholds increase gradually depending on the extent of inactivation of area MSTd[22]. Unfortunately, those results do not distinguish the two models, as there is no qualitative difference between the model predictions for partial MSTd inactivation (**Figure 7c**, red). Future experiments involving graded inactivation of VIP should be able to distinguish between the models due to the stark difference in their behavioural predictions.

The decoding strategies implied by the two models also have different consequences for how CCs should change during inactivation experiments (**Methods M10**). According to the extensive information model, VIP and MSTd are nearly independent, and both are decoded, so inactivating either area must scale up neuronal CCs in the other area (**Figure 7d – top**). In the limited information model, inactivating either area produces no significant changes in the other’s CCs (**Figure 7d – bottom**). This effect has different origins for MSTd and VIP. Although inactivating MSTd confines the readout to VIP, it also eliminates the high-variance noise components that VIP shared with MSTd: these two effects approximately cancel leaving CCs in VIP essentially unaffected. The results of VIP inactivation are simpler to understand: CCs in MSTd do not change much because VIP has little influence on behaviour to begin with.

## Discussion

Several recent experiments show that silencing brain areas with high decision-related activity does not necessarily affect decision-making[16–19]. To explain these puzzling results, we have developed a general, unified decoding framework to synthesize outcomes of experiments that measure decision-related activity in individual neurons and those that measure behavioural effects of inactivating entire brain areas. We know from the influential work of Haefner et al[14] how the behavioural impact (*readout weights*) of single neurons relates to their decision-related activity (*choice correlations*) in a standard feedforward network. We built on this theoretical foundation by adding three new elements that helped us relate the influence of multiple brain areas to both the magnitude of choice correlations, and the behavioural effects of inactivating those areas.

First, we have generalised their readout scheme to include multiple correlated brain areas by formulating the output of the decoder as a weighted sum of estimates derived from decoding responses of individual areas. In this scheme, the weight scales of individual estimates can be readily identified as the scaling of neuronal weights in the corresponding areas, providing a way to quantify the relative contribution of different brain areas. Second, we *postulated* that readout weights are mostly confined to a low-dimensional subspace of neural response that carries the highest response covariance, in both the extensive and limited information models. This postulate was instrumental to developing a theory of decoding that focused on the relationship between the overall scales of choice-related activity and neuronal weights, in lieu of their fine structures. Besides its mathematical simplicity, the resulting coarse-grained formulation confers an important practical advantage in that we can apply it without precisely knowing the fine structure of response covariance. Third, we used a straight-forward relation between behavioural threshold and the variance of the decoder to explicitly link the relative scaling of weights across areas to the behavioural effects of inactivating them.

Our theoretical result linking the behavioural influence of brain areas to their CCs and inactivation effects (**Equation 3.1 and 3.2**) is applicable only when neuronal weights within each area are mostly confined to the leading dimension of their response covariance. Although this requirement looks stringent, it is needed to explain the high CCs seen in experiments[15]. This claim might appear to be at odds with the fact that some earlier studies successfully predicted CCs that plateaued close to experimental levels using pooling models that did not explicitly take care of the above confinement[6,9]. However a closer examination revealed that these studies used a scheme in which decision was based on the average response of neuronal pools that were all uniformly correlated, a combination of model assumptions that in fact satisfies our requirement. Similar explanations apply to other simulation studies that used support-vector machines or alternative schemes that inadvertently restricted decoding weights to low-frequency modes of population response where shared variability was highest[12,30]. Thus our postulate is fully compatible with earlier work and in fact points to a more general class of models that can be used to describe the magnitude of CCs in those data.

Recent experiments show that reversibly inactivating area VIP in macaque monkeys does not impair animals’ heading perception, despite the fact that responses of VIP neurons are strongly predictive of perceptual decisions[18,21]. In contrast, inactivating MSTd does adversely affect behaviour even though MSTd neurons exhibit much weaker correlations with choice[22,23]. Assuming that both areas contribute to decision, we used our framework to infer decoding strategies that could account for these experimental results. Surprisingly, the data were consistent with two different schemes – *overweighting* or *underweighting* of VIP – depending on whether information was *extensive* or *limited*. A major implication of the finding from the extensive information model is that if a causal test of function (e.g., inactivation) reveals no impairments, it does not disprove that a brain area contributes to a task. The limited information model on the other hand suggests that area VIP is indeed of very little use to heading perception. In spite of this difference, both models share a basic attribute, namely, that decoding is suboptimal (although to very different extents, as discussed in the next section). Therefore our analysis reveals that the observed discrepancy between decision-related activity and effects of inactivation is not peculiar, and is actually expected from systems that integrate information across brain areas in a suboptimal fashion. The nature of this suboptimality can be understood intuitively by drawing an analogy to cue combination. Imagine there are two cues *x* and *y*, and you use a suboptimal strategy in which a larger weight is allocated to the less reliable cue *y*. If *y* is removed thereby forcing you to rely completely on *x*, then your behavioural precision might not change very much if the reduction in information from losing *y* is offset by the gain in information from *x*. On the other hand, if you mostly ignored *y* to begin with, then once again you will be unaffected by its removal. Either “too much” or “too little” weighting of a brain area can lead to suboptimal performance, both in a way that leaves the behavioural threshold largely unaltered following complete inactivation of that area.

## Decoding is suboptimal, but just how bad?

Although both models were suboptimal to some degree, the overwhelming distinction between them is the efficiency they imply for neural computation, where efficiency is the ratio of decoded information to available information. The efficiency of the limited information model is around 80%, independent of population size *N*. In contrast, the extensive information model encodes information that grows with *N*, while decoding is restricted to the least informative dimensions of neural responses. These decoders extract only a tiny fraction of the available information, resulting in an efficiency that falls inversely with *N*. For a modest-sized population of 1000 neurons, the efficiency is already less than 1%. Thus, the conventional model of correlated noise (with extensive information) is radically suboptimal, whereas the limited information model extracts an impressive fraction of what is possible, limited largely by noise.

It has previously been argued that the key factor that limits behavioural performance in complex tasks is suboptimal processing, not noise[38]. However, in simple tasks involving binary choices, and in areas in which most of the available information can be linearly decoded, it is unclear why the behaviour of highly trained animals should be so severely undermined by suboptimality. Moreover, radical suboptimality of the kind described here for the extensive information model implies tremendous potential for learning, as the neural circuits can continually optimize the computation by tuning the readout to more informative dimensions. This is hard to reconcile with the observation that behavioural thresholds in a variety of perceptual tasks typically saturate within a few weeks of training in both humans and monkeys[29,39–41]. In the presence of information-limiting noise, however, learning can only do so much, and performance must saturate at or below the ideal performance. Therefore we regard the limited information model as a much more likely explanation of our data, for otherwise one would need to posit that cortical computations discard the vast majority of available information. Note that suboptimal cortical computation might still account for information loss in the limited information model, as opposed to neural noise[38], but this information loss is now much more modest, probably around 20%.

A direct way to tell the two models apart would be to measure the structure of noise correlations. Unfortunately, this is not straightforward, because the differences between noise models giving extensive or limited information can be quite subtle[20]. In fact, there can be a whole spectrum of subtly different noise models with different information contents, lying between the two models that we have considered here. Therefore, a more accurate technique to determine the information content (which, after all, is a major reason why we care about noise correlations) is simply to record from hundreds of neurons simultaneously, and then decode the stimulus. This will provide a lower bound on the information available in the neural population. One can then compare the resultant population thresholds with the behavioural threshold to determine how suboptimal the decoding needs to be to account for behaviour. Eventually, we expect this strategy will be successful, but it will require advances in recording technology to be viable in the target brain areas. Meanwhile, by examining the key properties of the decoding strategy implied by the two models, we identified distinct predictions that are testable without large-scale simultaneous recordings. Specifically, they involve fairly simple experiments such as graded inactivation of VIP, and measurement of CCs in either VIP or MSTd while the other area is inactivated (**Figure 7**). Future experiments will test each of these predictions to provide novel evidence about the information content and decoding strategy used by the brain.

## Limitations of the framework and possible extensions

Similar efforts to deal with outcomes of correlational and causal studies using a coherent framework are rarely undertaken, despite their significance. To our knowledge, there is only one instance where this has been attempted before[42]. In that work, the authors used a recurrent network model with mutual inhibition between populations[43,44] to reconcile choice-related activity and the effect of silencing neurons. Although their study was similar to ours in spirit, their goal was different. They showed that inactivation just before a decision, when activity was highly correlated with the choice, had less impact on the behaviour than inactivation near the stimulus onset. This addresses a *temporal*, as opposed to a *spatial*, dissociation between correlation and causation, so a model with recurrent connectivity was essential to explain their findings. In contrast, we wanted to account for the discrepancies between measures of correlation and causation across brain areas. This latter phenomenon is entirely within the realm of standard feedforward network models in which both populations causally contribute, rather than compete to drive behaviour, and differ only in terms of the relative strength of their contributions.

Time-varying weights have been shown to better predict animals’ choice in certain tasks[45], and psychophysical kernels are sometimes skewed towards one end of the trial[46,47], suggesting that decoding could also be suboptimal in time. Such temporal weighting of information would naturally arise from recurrent connectivity, which is beyond the scope of this work. But it can also originate in feedforward networks, possibly through a gating mechanism that blocks the integration of neural responses beyond a certain time.[32]

Other studies have considered that choice-related activity might arise from decision feedback[46,48,49]. Indeed, pure decision feedback to an area would create apparent sensitivity to sensory signals, even in the absence of direct feedforward input to the target neurons[46,48,49]. In such a case, neural sensitivity to the stimulus would then be precisely equal to the animal’s sensitivity. In the absence of other sources of variability, response fluctuations would be perfectly correlated with fluctuations in the fed-back choice, producing choice correlations of 1. Of course there would be additional variability in the neural responses, and this would dilute both the choice correlations and neural tuning by equal amounts, giving rise to measured CCs that should match the optimal CCs (**Equation 2.1**). Even if there are other feedforward sensory components to the neural responses, direct decision feedback will pull the choice correlations toward this optimal prediction. Thus, simple decision feedback cannot account for the pattern of CCs observed in our VIP data, which are two to three times larger than predicted from optimal inference or direct decision feedback (**Figure 3**). Conversely, as we demonstrated through supplementary modeling, adding feedback or recurrent connections may not affect the suboptimal readout weights inferred using our scheme, even when those connections modulate responses along the decoded dimensions (**Figure S15**). Nevertheless, future expansions of our work should account for more general recurrent connectivity to study how neural circuits simultaneously integrate information across space and time. In particular, recurrent networks also include decision feedback as a special case, and might help test alternative theories on the origins of choice correlations[1,46].

Finally, while VIP inactivation did not impair heading discrimination, MSTd inactivation partially impaired the animal’s ability to perform the task. The fact that MSTd inactivation did not completely abolish performance cannot be accounted for by our two-population models unless the inactivation was only partial and/or VIP is read out to some degree. Additionally, we cannot exclude the possibility that VIP is merely correlated with behaviour and that a third brain area besides MSTd contributes some task-relevant information. In fact, both of our models actually predict a somewhat bigger deficit following MSTd inactivation (**Figure 5c, 6a**) than is observed experimentally (**Figure 1b**). This highlights the importance of ultimately extending coding models to include more than two brain areas.

As neuroscience moves towards ‘big data’, there is a greater need for theoretical frameworks that can help discern simple rules from complex multi-neuronal activity[50]. We believe our work responds to this challenge and, despite its limitations, takes us closer to bridging the brain-behaviour gap for binary-decision tasks.

## METHODS

### M1. Choice correlations in a linear feedforward model

Consider a standard feedforward decision process in which the neural response **r**~𝓝(**f**, Σ) is read out with weights **w** to generate an estimate 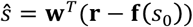. Choice correlations **C** in this scheme were previously shown[14,15] to be related to neuronal weights and response covariance according to 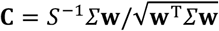 where 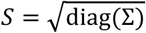. We can decompose these choice correlations into a sum of components arising from the individual noise modes of the *N*x*N* covariance matrix Σ as: 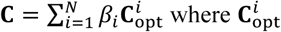 is the component of choice correlations generated from noise fluctuations along the *i^th^* mode when decoding weights **w** are optimal (**Supplementary notes S1, S2**). 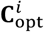 depends on the shape of the *i*^*th*^ noise mode **U**^*i*^, the amplitude of the signal **f**′ (the derivative of the neurons’ tuning curves), and the optimal threshold ϑ according to:

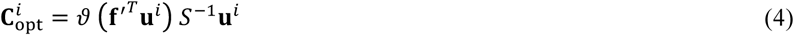

If decoding is optimal, then multipliers *β_i_*≡1 so the choice correlation *C_k,opt_* of neuron *k* becomes 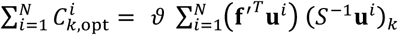 which reduces to *ϑ*/*ϑ* (**Supplementary note S2**) in agreement with earlier work[15]. In general however, multipliers *β_i_* will be different from 1 and can be estimated by regressing measured choice correlations **C** against the corresponding component 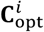

### M2. Weight scaling factors for unbiased decoding

Let **w**=(*a_x_***w**_1_, *a_y_***w**_y_)^*T*^ denote the readout weights of neurons where *a_x_* and *a_y_* represent the scaling of weights in the two populations *x* and *y*. To ensure unbiased decoding both before and after inactivation of the individual populations *x* or *y*, 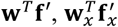, and 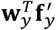 must all be equal to 1 where 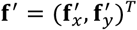 denotes the derivatives of the tuning curves of neurons in *x* and *y* (**Supplementary note S0**). This yields the constraint that *a_x_ + a_y_* = 1 at all times.

### M3. Relation between behavioural threshold and weight scaling factors

Behavioural threshold *ϑ* is proportional to the square root of the decoder variance (with proportionality of 1 for threshold of 68% correct), so ϑ^2^ = **w**^*T*^ Σ**w**. If decoding is confined to the subspace of leading eigenmodes of Σ spanned by neurons within *x* and *y* (**u**^*x*^ and **u**^*y*^), then **w**_*x*_ α *a_x_***u***x* and **w**_*y*_ α *a_y_**u**^y^* where the constants of proportionality are chosen to ensure unbiased decoding. In this case, the behavioural threshold can be expressed purely in terms of weight scaling factors and the variance originating from noise within the noise modes as (**Supplementary note S4**):

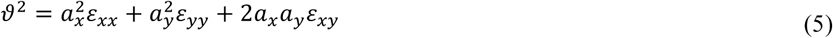

where *ε*_*xx*_ and *ε*_*yy*_ are the magnitudes of noise within *x* and *y*, and *ε*_*xy*_ is the magnitude of correlated noise. Thresholds following inactivation can be determined by setting the weight scaling factor for the inactivated area to zero, yielding 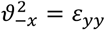 and 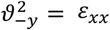.

### M4. Subjects and Behavioural Task

Six adult rhesus monkeys (A, B, C, J, S, U, and X) took part in various aspects of the experiments. Three animals were employed in each of the MSTd (C, J and S) and VIP (X, B and J) inactivation experiments. Two animals provided the neural data from each brain area (A and C for MSTd; C and U for VIP). All surgical and experimental procedures were approved by the Institutional Animal Care and Use Committees at Washington University and Baylor College of Medicine, and were performed in accordance with institutional and NIH guidelines. All animals were trained to perform a heading discrimination task around psychophysical threshold. In each trial, the subject experienced a real or simulated forward motion with a small leftward or rightward component (angle *s*, **Figure 1a**). Subjects were required to maintain fixation within a 2×2◦ electronic window around a head-fixed visual target located at the center of the display screen. At the end of each 2-s trial, the fixation spot disappeared, two choice targets appeared and the subject made a saccade to one of the targets to report his perceived heading relative to straight ahead. Nine logarithmically spaced heading angles were tested (0◦, ±0.5◦, ±1.3◦, ±3.5◦, and ±9◦ for monkeys A and J, 0◦, ±1◦, ±2.5◦, ±6.4◦, and ±16◦ for monkeys B, C, S and U), including the ambiguous case of straight ahead motion (*s* = 0◦). These values were chosen to obtain near-maximal psychophysical performance while allowing neuronal sensitivity to be estimated reliably for most neurons[21,23]. Subjects received a juice reward for indicating the correct choice. For trials in which the ambiguous heading was presented, rewards were delivered randomly on half of the trials. The experiment consisted of three randomly-interleaved stimulus conditions (vestibular, visual, and combined). In the vestibular condition, the monkey was translated by a motion platform while fixating a head-fixed target on a blank screen. In the visual condition, the motion platform remained stationary while optic flow simulated the same range of headings. Under the combined condition, both inertial motion and optic flow were provided. Each of the 27 unique stimulus conditions (9 heading directions×3 cue conditions) was repeated at least 20 times, for a total of 540 discrimination trials per recording session. Identical stimuli and trial structure were employed during both neural recordings and inactivation experiments.

### M5. Neural recordings

Activity of single neurons in areas MSTd and VIP was recorded extracellularly using epoxy-coated tungsten microelectrodes (impedance of 1–2 MΩ). Area MSTd was located using a combination of magnetic resonance imaging (MRI) scans, stereotaxic coordinates (~15 mm lateral and ~3–6 mm posterior to AP-0), white/gray matter transitions, and physiological response properties. In some penetrations, electrodes were further advanced into the retinotopically organized area MT[23]. Most recordings concentrated on the posterior/medial portions of MSTd, corresponding to more eccentric, lower hemifield receptive fields in the underlying area MT. To localize area VIP, we first identified the medial tip of the intraparietal sulcus and then moved laterally until there was no longer directionally selective visual response in the multiunit activity, as described in detail previously[21].

### M6. Estimation of Behavioural and Neuronal thresholds

Behavioural performance was quantified by plotting the proportion of ‘rightward’ choices as a function of heading (the azimuth angle of translation relative to straight ahead). Psychometric data were fit with a cumulative Gaussian function with mean μ and standard deviation *ϑ*, and this standard deviation defined the psychophysical threshold, corresponding to 68% correct performance (*d*^′^=1, assuming no bias, i.e. μ=0°).

For the analysis of neuronal responses, we used the linear Fisher information *J* which is simply a measure of the signal-to-noise ratio: signal power divided by noise power. The linear Fisher Information captures all of the Fisher information in responses generated from the exponential family with linear sufficient statistics. Its inverse is exactly equal to the variance of an unbiased, locally optimal linear estimator (for differentiable tuning curves and nonsingular noise covariance). We defined the square root of this variance (i.e. the standard deviation of the estimator) to be the neuronal discrimination threshold, which corresponds to 68% accuracy in binary discrimination. This threshold can be obtained directly from the neuron’s tuning curve and noise variance as follows:

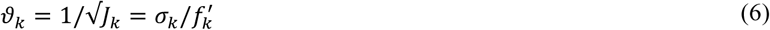

where *ϑ_k_* and *J_k_* are the threshold and linear Fisher information[51] for neuron *k*, 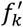 is the derivative of the neuron’s tuning curve at the reference stimulus (0◦), and 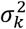 is the variance of the neuronal response for that stimulus. Neuronal thresholds computed using the above definition were very similar to those computed using a traditional approach based on neurometric functions constructed from the responses of the recorded neuron and a presumed ’antineuron’ with opposite tuning[52] (**Supplementary Figure 3**).

### M7. Estimation of Choice correlation

To quantify the relationship between neural responses and the monkey’s perceptual decisions, we first computed choice probabilities (CP) using ROC analysis[53]. For each heading, neural responses were sorted into two groups based on the choice that the animal made at the end of each trial. In previous studies, the two choice groups were typically related to the preferred and non-preferred stimuli for a given neuron[21,23]. In this study, in order to appropriately compare different neurons in a population code, the two choice groups were simply rightward and leftward choices; hence, CPs may be greater than or less than 1/2. ROC values were calculated from these response distributions, yielding a CP for each heading, as long as the monkey made at least 3 choices in favor of each direction. To combine across different headings, we computed a grand CP for each neuron by balanced *z*-scoring of responses in different conditions, which combines *z*-scored response distributions in an unbiased manner across conditions, and then performed ROC analysis on that combined distribution[54]. The CPs were then converted to choice correlations according to 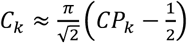 (refs. [14,15]) where *CP_k_* and *C_k_* are the choice probability and choice correlation of neuron *k* respectively (**Supplementary note S0**). Due to the convention we chose for computing CPs, the resulting choice correlation could be positive or negative depending whether a neuron predicted *rightward* choices by increasing or decreasing its response relative to reference stimulus. For an optimal decoder, the sign of a neuron’s choice correlation should match the sign of the derivative of its tuning curve, so we modified the definition of ref.[15] **(Equation 2.1)** to accommodate our sign convention, yielding 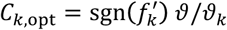 where sgn denotes the signum function.

There were neurons in both MSTd and VIP whose choice-related activity during the visual condition is anticorrelated with their signal-related activity[21,23]. Further analysis showed that heading preferences of these neurons during visual and vestibular conditions differed. Therefore the analysis of data collected during the visual condition presented in the Supplementary notes included only the subset of recorded neurons that had similar heading preferences as in the vestibular condition[23] (MSTd: 66/129 neurons; VIP: 63/88 neurons).

### M8. Noise covariance of extensive information model

Pairwise neuronal recordings carried out separately in areas VIP and MSTd were used to estimate noise correlations between pairs of neurons, *R_ij_* = Corr(*r_i_, r_j_|s* = 0), where *r_i_* and *r_j_* are the responses of neurons *i* and *j*, and correlation coefficients were computed by averaging over trials with headings near 0°. The same recordings were used to compute signal correlations, 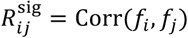, where *f_i_* and *f_j_* are the tuning curves of neurons *i* and *j*, and the correlation coefficients were computed by averaging over a uniform distribution of headings in the horizontal plane. The typical noise correlations, 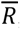, were then modeled as linearly proportional to the signal correlations:

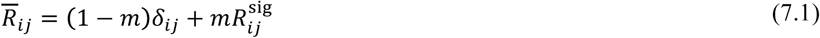

where *δ*_*ij*_ is the Kronecker delta function (*δ*_*ij*_ is 1 when *i* = *j*, and 0 otherwise) and *m* is the slope of the relationship between signal correlations and noise correlations. This slope was much steeper in VIP than MSTd[21]. For the vestibular condition, slopes were found to be *m_M_* =0.19±0.08 and *m_v_* =0.70±0.16 within MSTd and VIP respectively, and for the visual condition they were *m_M_* =0.12±0.09 and *m_v_* =0.50±0.14. The above fits determined the average relationship between noise and signal correlations, but there was considerable diversity around this trend. To emulate this diversity, we used a technique similar to the one proposed in ref. [31]. Specifically, we sampled correlation coefficient matrices 𝑅 from a Wishart distribution with a mean matrix 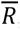 given by **equation 7.1** and the fitted slope *m*, and rescaled them to ensure *R_ii_* = 1. The number of degrees of freedom for the Wishart distribution was adjusted so sampled matrices had the same uncertainty in slope *m* as the data when subjected to the same fitting procedure. Covariance matrices were generated by scaling the correlation coefficients by the standard deviations for each neuron. Model variances were set equal to the mean responses, so the standard deviation of neuron *i* is 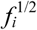. Thus the covariance Σ is related to correlation coefficients *R* by 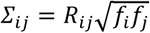. Correlations between responses of MSTd and VIP neurons were not measured experimentally, so the slope *m_MV_* of any linear trend relating noise and signal correlations between the two areas was not known. We explored different possibilities by varying *m_MV_* according to:

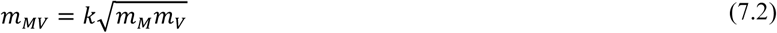

where *k*ϵ[0,1). Each value of *k* produced correlation between areas with magnitude *ε_MV_* which was expressed as *ε_MV_* = *γ ε*_*MM*_.

### M9. Noise covariance of limited information model

If the information reaching MSTd (*M*) and VIP (*V*) is not perfectly redundant across the populations, then the resulting covariance matrix will be of the form:

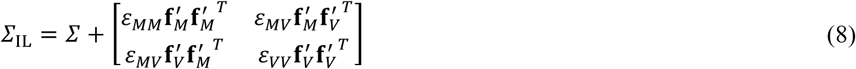

where 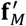 and 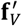 are derivatives of tuning curves of the neurons in *M* and *V* respectively, and 𝛴 is the noise used in the extensive information model. Whereas 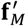 and 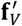 can be estimated by measuring the tuning curves of individual neurons, precisely estimating *ε*_*MM*_, *ε_VV_*, and *ε*_*MV*_ is difficult even with large-scale recordings as their magnitudes may be very small compared to the magnitude of noise in Σ. Nevertheless, we know that for large populations, the behavioural threshold will be dominated by the magnitude of information-limiting correlations. Specifically, they are related through the relative scaling of decoding weights in **equation 5** where *M* and *V* take the places of *x* and *y*. Consequently, we can determine *ε*_*MM*_ and *ε_VV_* from behavioural thresholds following inactivation using 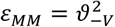 and 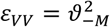. We can then use **equation 5** in conjunction with **equation 3.2** to determine both the ratio *a_M_*/*a_V_* of weight scalings and the magnitude of correlation between populations *ε_MV_*=*γε_MM_*.

### M10. Effects of inactivation on choice correlations

Complete inactivation of one of the areas will affect neuronal choice correlations in the non-inactivated area. If **C**_*x*_ and 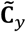 denote the choice correlations of neurons in area *x* before and after inactivation of *y*, then it can be shown that 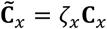 and similarly 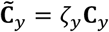 where scalars ζ_*y*_ and ζ_*y*_ are (**Supplementary note S10**):

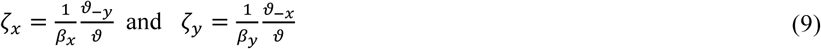

where *β_x_* and *β_y_* are the multipliers that relate the observed and optimal patterns of neuronal choice correlations in areas *x* and *y*. The above equation implies that choice correlations in the active area will increase by a factor proportional to the behavioural effect of inactivating the other area. Intuitively, this is because inactivating an area that was very important for behaviour will dramatically increase the burden on the active area, leading to an increase in the magnitude of choice-related activity.

## Acknowledgements

The work was supported by NIH R01 DC04260, R21 DC014518 and the Simons Collaboration for the Global Brain, grant #324143. A.P. was supported by a grant from Simons Global Brain Initiative and the Swiss National Foundation (#31003A_143707). We thank Adam Zaidel, Yong Gu, & Aihua Chen for performing the neural recordings, as well as Sheng Liu & Yong Gu for performing the muscimol inactivation experiments.

